# Groundwater metabolome responds to recharge in fractured sedimentary strata

**DOI:** 10.1101/2022.03.17.484695

**Authors:** Christian Zerfaß, Robert Lehmann, Nico Ueberschaar, Carlos Sanchez-Arcos, Kai-Uwe Totsche, Georg Pohnert

**Affiliations:** Department of Bioorganic Analytics, Institute of Inorganic and Analytical Chemistry, Friedrich Schiller University, Jena, Germany; Department of Hydrogeology, Institute of Geoscience, Friedrich Schiller University, Jena, Germany; Mass Spectrometry Platform, Faculty for Chemistry and Earth Sciences, Friedrich Schiller University, Jena, Germany

**Keywords:** Groundwater, dissolved organic matter, DOM, recharge, discharge, metabolomics, aquifer, fractured sedimentary strata, triassic limestone-mudstone

## Abstract

Understanding the sources, structure and fate of dissolved organic matter (DOM) in groundwater is paramount for protection and sustainable use of this vital resource. On its passage through the Critical Zone, DOM is subject to biogeochemical conversions. Therefore, it carries valuable cross-habitat information for monitoring and predicting the stability of groundwater ecosystem services and assessing these ecosystems’ response to fluctuations caused by external impacts such as climatic extremes. Challenges arise from insufficient knowledge on groundwater metabolite composition and dynamics due to a lack of consistent analytical approaches for long-term monitoring. Our study establishes groundwater metabolomics to decipher the complex biogeochemical transport and conversion of DOM. We explore fractured sedimentary bedrock along a hillslope recharge area by a 5-year untargeted metabolomics monitoring of oxic perched and anoxic phreatic groundwater. A summer with extremely high temperatures and low precipitation was included in the monitoring. Water was accessed by a sampling well-transect and regularly collected for liquid chromatography-mass spectrometry (LC-MS) investigation. Dimension reduction of the resulting complex data set by principal component analysis revealed that metabolome dissimilarities between distant wells coincide with transient cross-stratal flow indicated by groundwater levels. Time series of the groundwater metabolome data provides detailed insights into subsurface responses to recharge dynamics. We demonstrate that dissimilarity variability between groundwater bodies with contrasting aquifer properties coincides with recharge dynamics. This includes groundwater high- and lowstands as well as recharge and recession phases. Our monitoring approach allows to survey groundwater ecosystems even under extreme conditions. The metabolome was otherwise highly variable lacking seasonal patterns and did not segregate by geographic location of sampling wells, thus ruling out vegetation or (agricultural) land use as primary driving factor. Patterns that emerge from metabolomics monitoring give insight into subsurface ecosystem functioning and water quality evolution, essential for sustainable groundwater use and climate change-adapted management.

## Introduction

The recent IPCC report highlights the potential impact of climate change on groundwater quality and quantity. It though notes a lack of quantitative data on groundwater development that prevents the preditction of potential impacts reliably (Caretta et al., 2022). Particularly the composition of dissolved organic matter (DOM) and its fate in groundwater reservoirs are rather obscure. DOM signatures are essential for assessing groundwater ecosystem functioning and vulnerability to land use, water resource exploitation, and climate change-driven threats (Drake et al., 2020; Humphrey et al., 2016; Phillips et al., 2012; Riedel & Weber, 2020; Rodell et al., 2018; Rodríguez-Cardona et al., 2022; United Nations Office for Disaster Risk Reduction (UNDRR), 2021; Wada et al., 2012). Groundwater DOM sources, composition, persistence, and transformation reflect biogeochemical processes below-ground (Kaiser & Kalbitz, 2012; J. Lehmann & Kleber, 2015; Li et al., 2017; Roth et al., 2019; Schmidt et al., 2011). DOM also plays a crucial role in the transport of organics and in the supply of the below-ground microbial communities. These multiple functions attract a high interest for DOM in Critical Zone (CZ) research worldwide (Grant & Dietrich, 2017; Küsel et al., 2016; Li et al., 2017; Richardson, 2017; Riebe et al., 2017).

For sustainable resource management and groundwater protection, a sound understanding of recharge areas’ hydrogeological and ecological functioning is required (R. Lehmann & Totsche, 2020). Multi-disciplinary, temporally highly-resolved and long-term monitoring efforts can provide valuable insight and support concepts for management. However, such approaches are still rare, except for targeted investigations of contaminants (Fernandes et al., 2021). To date, we have no information about spatial and temporal patterns and dynamics of the entirety of metabolites and their transformation products (here termed as metabolome) in the groundwater.

Such information is especially required for vulnerable recharge areas where cross-compartment matter exchange (R. Lehmann & Totsche, 2020) and dynamics of microbial activity contribute to the evolution of groundwater quality (Benk et al., 2019). Fractured sedimentary bedrock with mixed lithologies hosts important groundwater reservoirs worldwide. These reservoirs are challenging to investigate due to the interplay of transient flow patterns and various sedimentary or surface-fed sources of organic matter. In addition, the dynamics of microbial communities, their metabolic activity, and their longterm persistence influence the groundwater metabolome. Assessing the temporal dynamics of the groundwater metabolome may provide cross-habitat information in multiple ways: Metabolic composition reflects the input from above ground and contributions from mobile and rock-associated microbial populations in high-permeability (fractures) and low-permeability (rock matrices) environments. In addition, abiotic transformations contribute to metabolic diversity below ground (Chorover & Amistadi, 2001).

Untargeted metabolomics techniques are emerging tools in environmental monitoring of compound transportation and transformation (Crimmins & Holsen, 2019). They have also been successfully used to characterize DOM in ground- (Aukes & Schiff, 2021; Benk et al., 2019; Hofmann et al., 2020), soil-(Danczak et al., 2020; Lu et al., 2020; Seifert et al., 2016; Ueberschaar et al., 2021) and surface-waters (Aukes & Schiff, 2021; Danczak et al., 2020; Drake et al., 2020; Garayburu-Caruso et al., 2020; Hawkes et al., 2020; Lu et al., 2020; Lynch et al., 2019; Patriarca & Hawkes, 2021; Pemberton et al., 2020; Raeke et al., 2017; Seifert et al., 2016; Tanentzap et al., 2019; Wologo et al., 2021). As sub-surface metabolites are not well characterized, exact compound annotation is challenging. Therefore, previous studies focused on selected marker molecules and compound class categorization. However, fingerprint analyzes by multivariate approaches such as principal component analysis also proved a powerful tool for the analysis of aggregated monitoring data (Benk et al., 2018, 2019; Schwab et al., 2017; Ueberschaar et al., 2021). These approaches allowed for categorisation of similarities across compartments that can be linked to subsurface connectivity (Küsel et al., 2016; R. Lehmann & Totsche, 2020) and the composition of groundwater microbiomes (Benk et al., 2019; Yan et al., 2020).

Here we present a long-term, high-resolution analysis of groundwater metabolome patterns in Triassic limestone-mudstone alternations of a topographic recharge area (Figure 1). We profiled both perched and phreatic groundwater for over 5 years at multiple locations. The assessment is based on untargeted metabolomics using data from liquid chromatography coupled to mass spectrometry (LC-MS). A two-step data analysis approach is chosen. Metabolome data per individual monitoring well is analyzed over time giving seasonal and intra-annual data at specific locations. In addition, data are analyzed by generating a dissimilarity score at each timepoint, by which evolving metabolomes are investigated over time. While a detailed analysis of the metabolomes at a single time point was conducted previously (Sanchez-Arcos et al., 2022), the data analysis presented here aimed at exploring the metabolome structure and dynamics in distinct zones of the groundwater recharge area shown in Figure 1.

**Figure 1.**
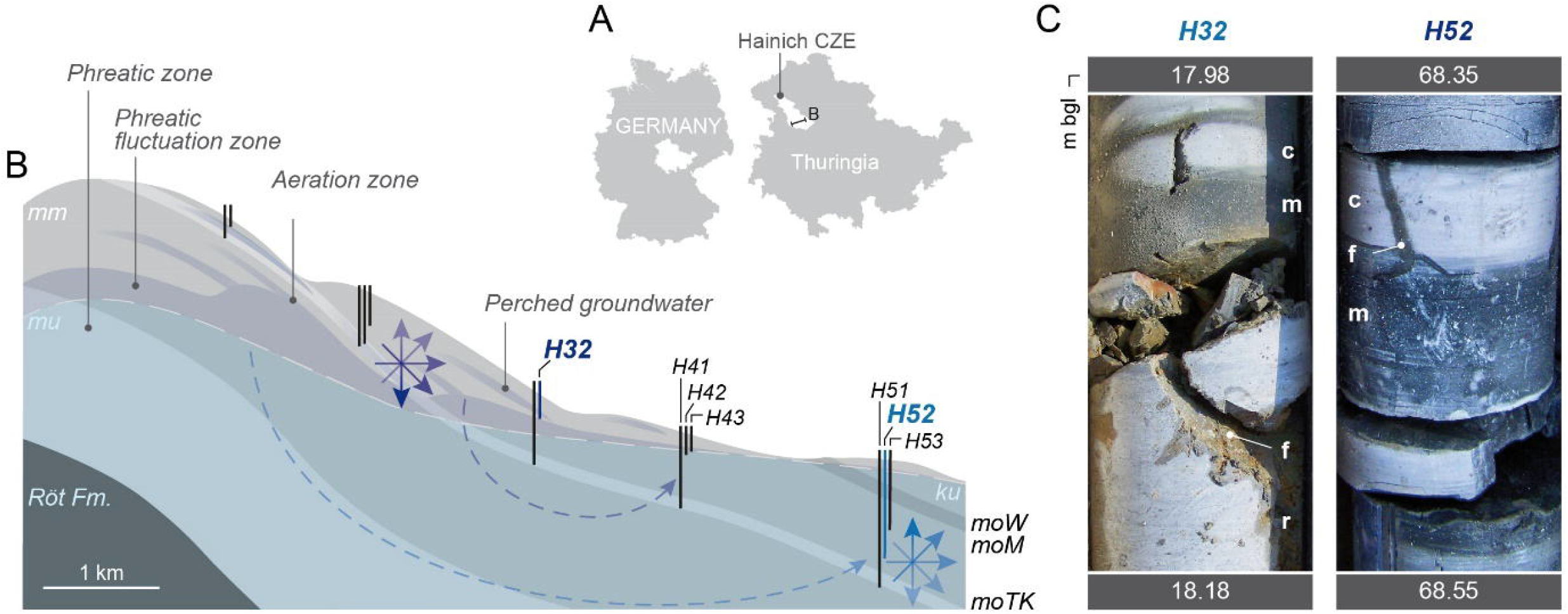
Configuration and aquifer rock properties of the study site. **(A)** Location of the well transect in central Germany. **(B)** Schematic cross section of the hillslope groundwater recharge area of the Hainich Critical Zone Exploratory (CZE). Arrow roses highlight transient and prevailing flow directions in two contrasting compartments, the deep aeration zone (downward/stratiform > upward) and the phreatic zone (upward > downward) in discharge direction. Dashed blue lines show presumed groundwater flow patterns. Bedrock strata belong to lithostratigraphic subgroups Lower Muschelkalk (mu), Middle Muschelkalk (mm), Upper Muschelkalk (mo) with Trochitenkalk formation (moTK), Warburg Fm. (moW), Meissner Fm. (moM), above the regional flow barrier (Röt formation, Upper Buntsandstein). **(C)** Examples of drill cores from the screen sections of the monitoring wells, representing contrasting compartments with similar lithology. Left: Weathered mudrock and stained fractures indicate oxic conditions in diverse aeration zone habitats. Right: Rock colors and brittle mudstone beds indicate oxygen-deficient conditions in low-weathered strata. c: calcimudstone, f: fracture, m: mudrock/-stone, r: rudstone (bioclastic limestone).

The metabolomic dataset is embedded in an extensive assessment of environmental data. This includes water levels, inorganic ions, DOC (sum parameter) and other basic hydrochemical parameters such as pH, temperature, dissolved oxygen and electric conductivity. These large sets of basic parameters allow for the reconstruction of flow regimes and ecosystem compartmentalization (R. Lehmann & Totsche, 2020). In this context, untargeted metabolomics enables the understanding of subsurface ecosystem functioning and functions by untangling the specific information contained in DOM. The metabolome reflects the signal resulting from the surface and subsurface sources (e.g. sedimentary carbon turnover) and all abiotic and biotic transformations of organic metabolites within the Critical Zone until the time of sampling. The groundwater metabolome thus contains cross-habitat information, potentially revealing whole-ecosystem functioning and stability. Notably, the metabolic output will not associate with individual organisms but with microbial communities. Thus, metabolomic assessment represents a proxy for metabolic activity in cross-habitat microbial populations.

## Materials and Methods

### Site description

The exemplary studied groundwater recharge area is part of the Hainich Critical Zone Exploratory (or observatory, CZO) in NW Thuringia, central Germany. It was established by the Collaborative Research Centre AquaDiva focussing on subsurface biodiversity, groundwater quality, and its surface- and subsurface controls (Küsel et al., 2016). The site represents fractured sedimentary bedrock environments, more specifically, widely distributed mixed carbonate-/siliciclastic rock alternations, as well as groundwater reservoirs in agricultural catchments with forested recharge areas in low-mountain terrain. The monitoring well transect spans ca. 7 km along the eastern hillslope of the Hainich mountain range, comprising different relief positions, aquifers, and depths (see Figure 1). The wells access perched groundwater bodies in the upper slope of a thick aeration zone, and phreatic groundwater in downslope positions with depths between 7 and 88 meters below ground level (Schwab et al., 2017). In this study, analysis includes data from each three wells in mid- to footslope positions (H41-H43 and H51-H53, phreatic zone), and one upper midslope well (H32, perched groundwater), representing the most continuous time series in our long-term metabolomic monitoring.

We demonstrate our metabolome analysis for two representative compartments within the studied groundwater recharge area: perched groundwater in the shallow aeration zone (well H32), and deeper phreatic groundwater (H52) that locates in between two other phreatic zone observer wells (H51 and H53).

### Groundwater sampling

Groundwater samples were provided during joint, usually monthly sampling campaigns of the CRC AquaDiva as described previously (Küsel et al., 2016; R. Lehmann & Totsche, 2020). Each 5 litres of groundwater per well was sampled in triplicate borosilicate bottles by employing submersible pumps (MP1 or SQE 5-70, Grundfos, Denmark) and PE-HD tubing, after representative conditions were achieved by monitoring physicochemical parameters until stabilisation (R. Lehmann & Totsche, 2020).

### Untargeted metabolomics by LC-MS

Samples collected during a three-day campaign were stored dark and chilled until subsequent transfer and immediate processing. Metabolite extraction was carried out by GF/C filtering (1.5 μm, VWR #516-0875) followed by solid phase extraction (SPE) on Strata-X 33 μm polymeric reversed phase cartridges (500 mg, Phenomenex #8B-S100-HCH). Prior to extraction, the SPE cartridges were placed on a Supelco Visiprep DL 24 manifold (Merck/Sigma-Aldrich #57265) and conditioned with 5 ml LC-MS grade methanol. Subsequently, GF/C filters placed in a Swinnex 47 mm diameter filter holder (Carl Roth, #XE88.1) were assembled onto the cartridges, and connected via Teflon tubes to the sample bottles. Flow of sample was maintained by applying vacuum (up to −500 mbar as measured by a manifold-attached manometer). After 5 l groundwater samples had run through, 5 ml of LC-MS grade water were applied for washing the SPE cartridges (GF/C filter already detached), and vacuum was applied for 5 minutes for cartridge drying. For elution, 4 ml of a 1:1 mixture methanol/acetonitrile (LC-MS grade) was applied and extracts were collected in 4 ml glass vials. The solvent was evaporated by applying a nitrogen stream or vacuum and subsequently dissolved in 100 μl of a 1:1 mixture of methanol/tetrahydrofuran (LC-MS grade). Samples were transferred to LC glass vials (with insets) and if sediments formed, clear supernatants were transferred to new vials by pipetting. From these, 1 μl was injected into the LC-MS in technical triplicates.

LC-MS was performed on a Dionex UltiMate 3000 chromatography system coupled to a Q-Exactive Plus orbitrap mass spectrometer (Thermo Fisher Scientific, Germany). A heated electrospray ionization (HESI) source was used with the following ionization and mass spectrometry settings: mass range 100-1,500 m/z, resolution 70,000 m/Δm, AGC target 3*106, polarity positive and negative (only positive used in later analysis), maximum inject time 200 ms, spray voltage 3.0 kV (positive mode) and 2.5 kV (negative mode), capillary temperature 360 °C. LC separation was conducted at 25 °C column temperature, with slightly different methods as follows: *LC Method 1 (samples until June 2017)*: LC-column: Kinetex Core Shell (Phenomenex; 1.7 u, C18, 100A, 100×2.1 mm). Solvent A: H2O, 2% acetonitrile, 0.1% formic acid. Solvent B: acetonitrile. LC-gradient: 0-0.2 min 100% A; 0.02-8 min linear gradient to 0% A; 8-9 min 0% A; 9.1-10 min 100% A. *LC Method 2 (samples March 2018 till July 2019)*: LC-column: Accucore TM C18 (Thermo Fisher Scientific, 2.1 x 100 mm, 2.6 mm). Solvent A: H2O, 2% acetonitrile, 0.1% formic acid. Solvent B: acetonitrile, 0.1% formic acid. LC-gradient: 0-0.2 min 100% A; 0.02-8 min linear gradient to 0% A; 8-12 min 0% A; 12.1-14 min 100% A. *LC Method 3 (samples from March 2020; equivalent gradient to LC Method 1 with longer recording towards the end of method):* LC-column: Kinetex Core Shell (Phenomenex; 1.7 u, C18, 100A, 100×2.1 mm). Solvent A: H2O, 2% acetonitrile, 0.1% formic acid. Solvent B: acetonitrile. LC-gradient: 0-0.2 min 100% A; 0.02-8 min linear gradient to 0% A; 8-11 min 0% A; 11.1-12 min 100% A.

### Data processing and statistical analysis

Data acquired by liquid chromatography coupled to mass spectrometry is accessible via the Metabolights repository (Haug et al., 2019) with study identifier MTBLS3450 (Zerfaß et al., 2021). Thermo Fisher Xcalibur raw LC-MS output files were converted to mzXML files with ProteoWizard (Chambers et al., 2012) and subsequently analyzed in R (R-CoreTeam, 2017) with the Bioconductor (Huber et al., 2015) package XCMS (Benton et al., 2010; Smith et al., 2006; Tautenhahn et al., 2008). Datasets were analyzed with standardized XCMS parameters, and crucial steps are described in the following. During data conversion to mzXML by ProteoWizard, positive-mode LC-MS traces were extracted in the time range 50-570 seconds, in which recordings were overlapping for LC-MS methods 1 to 3 as described above. The converted files were read into, and analyzed by, XCMS (function: readMSdata), peak identification and integration with parameters defined in centWaveParam, retention time adjustment was carried out with the obiwarp method, and grouping and peak filling from baseline data was executed with groupChromPeaks and fillChromPeaks, respectively). The table of integrated peaks was extracted with the featureValues function.

For hierarchical clustering of metabolite feature traces, peak intensities were z-normalized (Cooksey, 2020; Miller et al., 2015), and averages and standard deviations were calculated for the technical replicates (i.e. triplicate LCMS injections) and groundwater replicates (triplicates, standard deviation via pooled variance). Clustering was carried out by the R-function cor with Pearson correlation for complete observations. Variability was assessed by ANOVA with Benjamini Hochberg correction (Antonelli et al., 2019) (R-functions aov and p.adjust).

For principal component analyzes, zero peak intensities were changed to “not applicable” (NA), and the peak intensity table was log_2_-transformed. Again, averages and standard deviations were calculated for the technical replicates (i.e. triplicate LCMS injections) and groundwater replicates (triplicates, standard deviation via pooled variance). Principal component analysis (Ivanisevic & Want, 2019; Nguyen & Holmes, 2019; Worley & Powers, 2012) (PCA) was carried out with R’s prcomp function on centred peak intensity data, omitting NAs, on the groundwater-replicate averages dataset. With the resulting rotation data, sample averages and standard deviations were transformed into the PC1-PC2 space.

For long-term analysis of consistent analysis periods, data for one sampling point at different time points was fed into the PCA. Notably, measurement parameters may drift over time, such as for (a) changes in chromatography columns and LC methods and (b) due to the long time period, drifts in e.g. LC-column capacities and separation efficiency that may complicate integrated data-analysis involving retention time alignments of chromatograms. We therefore conducted an additional analysis where PCAs were carried out for each sampling campaign (i.e. time points) with a complete set of seven sampling locations from the Hainich CZE wells H32, H41-3, and H51-3 (compare Figure 1). Distances between each two datapoints (i.e. between the measurement averages) in the PC1-PC2 space were calculated, and the standard deviations of the distances were calculated via the pooled (weighted) variance in the direction of the connecting line as follows. Knowing the differences between datapoints in the direction of the PC1 and PC2 axis, respectively, the distance was inferred from Pythagoras’ law as

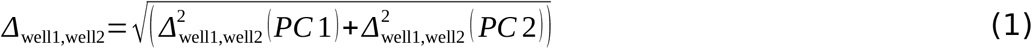

and the angle of the connecting line (measured from PC1 axis) was inferred by

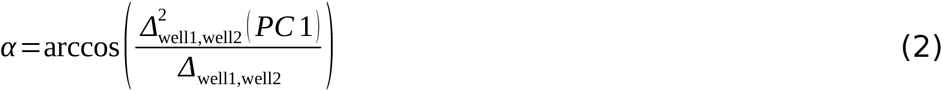

The standard deviation in the PC1-PC2 space was considered as elliptical space (Wang et al., 2015; Yuill, 1971) with limits defined by the respective standard deviations in the first two principal components (δ(PC1) and δ(PC2)). From these limits, the coordinates of the standard deviation ellipse for all angles α were associated with an angle φ on the limit circles, so that

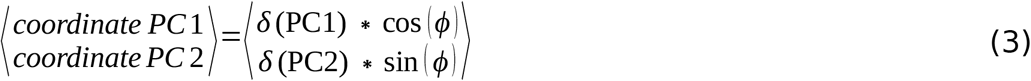

The length of the standard deviation vector at the angle α can therefore be calculated from the individual standard deviations δ(PC1) and δ(PC2) by determining the angle φ with the following equation

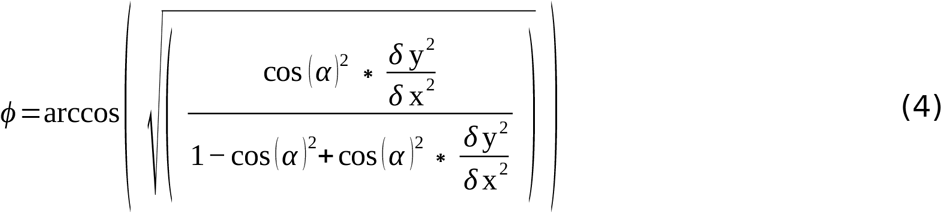

and using Pythagoras’ law as

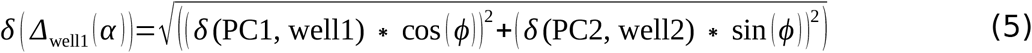

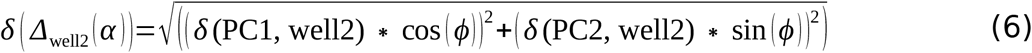

This allows to calculate a standard deviation for each of the sampling wells in the PC1-PC2 space, in the direction of the connecting line between the data points. The standard deviation of the well-distance was calculated via the pooled (weighted) variance from the individual standard deviations of the datapoints, with equal sample sizes (leaving the weighting term obsolete). The difference between the wells was thereafter divided by the standard deviation of the difference, which expresses the distance as multiples of its standard deviation:

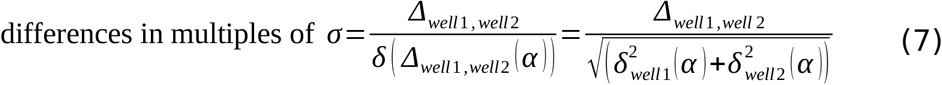

Note for statistical assessment that for a normal-distributed random sample, the 1σ and 2σ ranges mark the intervals expectedly containing 68% and 95% of observations, respectively (Wang et al., 2015). To analyze the dissimilarity of wells over time, the well differences in multiples of σ were compared by plotting them against the date of sampling campaigns.

## Results and Discussion

### Dimension reduction of metabolome data reveals responses to high-flow conditions in the aeration and phreatic zone

The long-term monitoring of groundwater bodies with contrasting aquifer properties along a topographic recharge area provided a metabolomic data set suitable for determining spatiotemporal patterns. Groundwater sampling, filtration, and extraction on reversed-phase material were optimized for reproducibility, recovery, and robustness. Extracts were measured with high resolution LC/MS using reversed-phase ultra high-pressure liquid chromatography (UHPLC) for separation. We focus here on pattern recognition in these data sets without the aim to elucidate the structure of the detected metabolites. The five-year monitoring dataset comprised 35 time points, when data was collected at three geographic locations from sampling wells at one (H32) or three (H41-H43, H51-H53) depths. This data set was systematically mined to identify patterns according to location, depth, season and recharge events. In addition, basic hydrochemical and conventional monitoring parameters, including groundwater levels and temperature were collected along with the metabolome. A systematic geographic analysis of the metabolomes from different wells at a single time point is reported elsewhere (Sanchez-Arcos et al., 2022).

The thick hillslope aeration zone, containing perched groundwater bodies (Figure 1) is accessed by well H32. Multiple pronounced periods of recharge in the winter half-year, interspersed by low-stand phases were observed there during the monitoring period from 2014 to 2020 (Figure 2 A). During recharge, the Ca/Mg-carbonate rich water is oxygenated (up to 3.8 mg/L dissolved oxygen, not shown) and the temperature dips (Figure 2 A). Further, nitrate is supplied via percolation in the winter half-year (Figure S1). Periods of low specific electrical conductivity during early recharge phases indicate short residence times (Figure S1). Dissolved organic carbon (DOC, sum parameter) did not show significant responses to recharge, whereas increasing sulphur/sulphate, sodium, calcium and decreasing potassium point to longer residence during recession phases (Figure S1) (Bakalowicz, 1994; Hudak, 2004).

**Figure 2.**
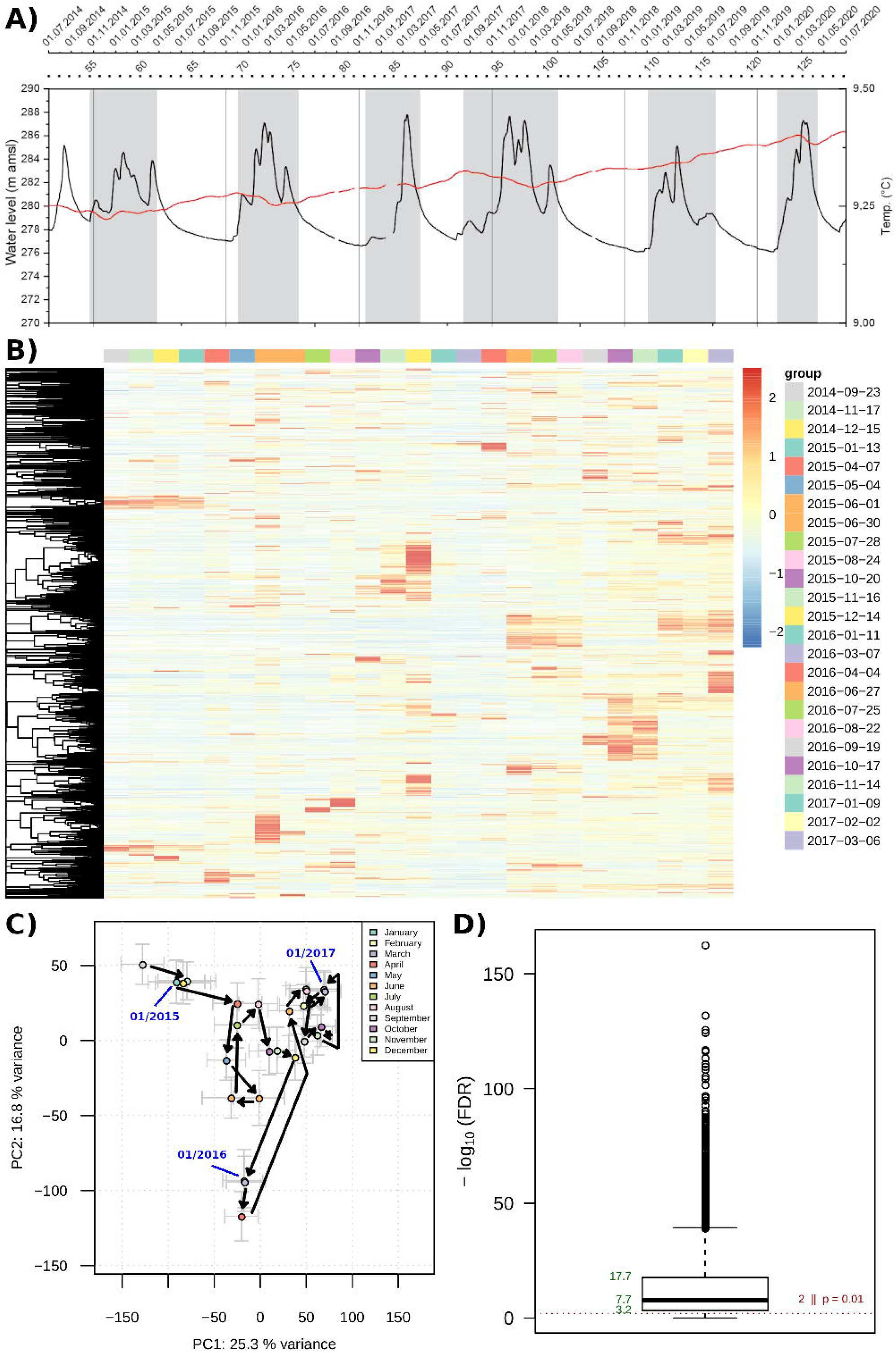
Single-well (H32) metabolome time-series analysis. **(A)** Groundwater level and temperature traces. Recharge phases (grey-shaded) are inferred from the multi-level head data and the environmental tracer temperature (red). Seasonal cooling is observed during the winter half-year highstands. **(B)** Hierarchical cluster analysis of z-normalised (Cooksey, 2020; Miller et al., 2015) metabolite feature intensities. **(C)** Principal component analysis of time-series data. Arrows are drawn as guides for chronological order of datapoints; the first datapoint of each year is labelled in blue font. **(D)** False discovery rate (FDR; from Benjamini-Hochberg corrected ANOVA) distribution as indicator of significance of metabolite feature intensity variability in the time series.

The metabolome of well H32 from 09/2014 to 03/2017 (consistent monitoring with mostly monthly sampling campaigns) revealed a highly variable composition, with more than 75% of the metabolite features significantly changing in intensity over time (Figure 2 B, D). Notably, the metabolomes at different time points were characterized by singular emergences of metabolite features, rather than recurrent profiles (Figure 2 B). The lack of recurrent signals may characterize the studied setting of thin-bedded alternations of mixed carbonate-/siliciclastic-rock. Lacking large fractures or karst conduits, such alternations represent retarded flow systems (White, 2012), thus generally exhibiting less pronounced DOM dynamics by input from the subsurface. DOM dynamics might result from multidirectional flow dynamics within groundwater recharge areas (R. Lehmann & Totsche, 2020), supplying water and DOM from different sources, including sedimentary carbon. Such sediments are common in freshwater aquifers worldwide, and the observed patterns might be prototypic (Franco et al., 2017).

Interannual variability may generally depend on weather and climatic conditions (droughts, precipitation, temperature) (Humphrey et al., 2016; Phillips et al., 2012; Riedel & Weber, 2020; Rodell et al., 2018; United Nations Office for Disaster Risk Reduction (UNDRR), 2021). Elsewhere, geochemical tracer investigations revealed that responses to intra-annual periodicity in precipitation and associated recharge is reflected in the groundwater composition (Bonneau et al., 2018; Bowen et al., 2019; Jasechko et al., 2014; Tweed et al., 2005). Emergence and dissipation of metabolite signals could be linked to these factors within the course of a year.

The first two (determining) principle components revealed considerable patterns when metabolic signatures throughout the sampling period were analyzed (Figure 2 C). The most striking was an observed shift coinciding with the high-flow period 12/2015-05/2016, causing a disturbance in the PCA (Figure 2 C, lower half). Similarly, the high-flow periods 11/2014-06/2015 (upper-left of PCA) and 11/2016-04/2017 (Figure 2 C, upper right) caused separate clusters, interspersed by distinct phases such as the low-stand phase 06/2015-11/2016 (Figure 2 C, center-left).

The much deeper phreatic zone well H52 (Figure 1) was characterized by damped, though high-magnitude water table fluctuation of more than 18 m due to seasonal recharge dynamics (Figure 3 A; Supplementary Figure S2). Compared to the perched groundwater, the oxygen-free Ca/Mg-carbonate water is low in nitrate and high in sulfate. As indicated by less surface-fed input, this aquifer zone is characterized by more isolated conditions (R. Lehmann & Totsche, 2020). Recharge phases can be recognized by ascending waters (Figure 3 A), supplying calcium and sulfate from laying strata, and causing decreased EC 25, with reduced sodium and potassium during high flow periods. These observations highlight considerable cross-stratal exchange (R. Lehmann & Totsche, 2020) in the stratified bedrock. DOC, again, showed insignificant fluctuation, pointing to the limited merit of the sum parameter in such a sedimentary setting.

**Figure 3.**
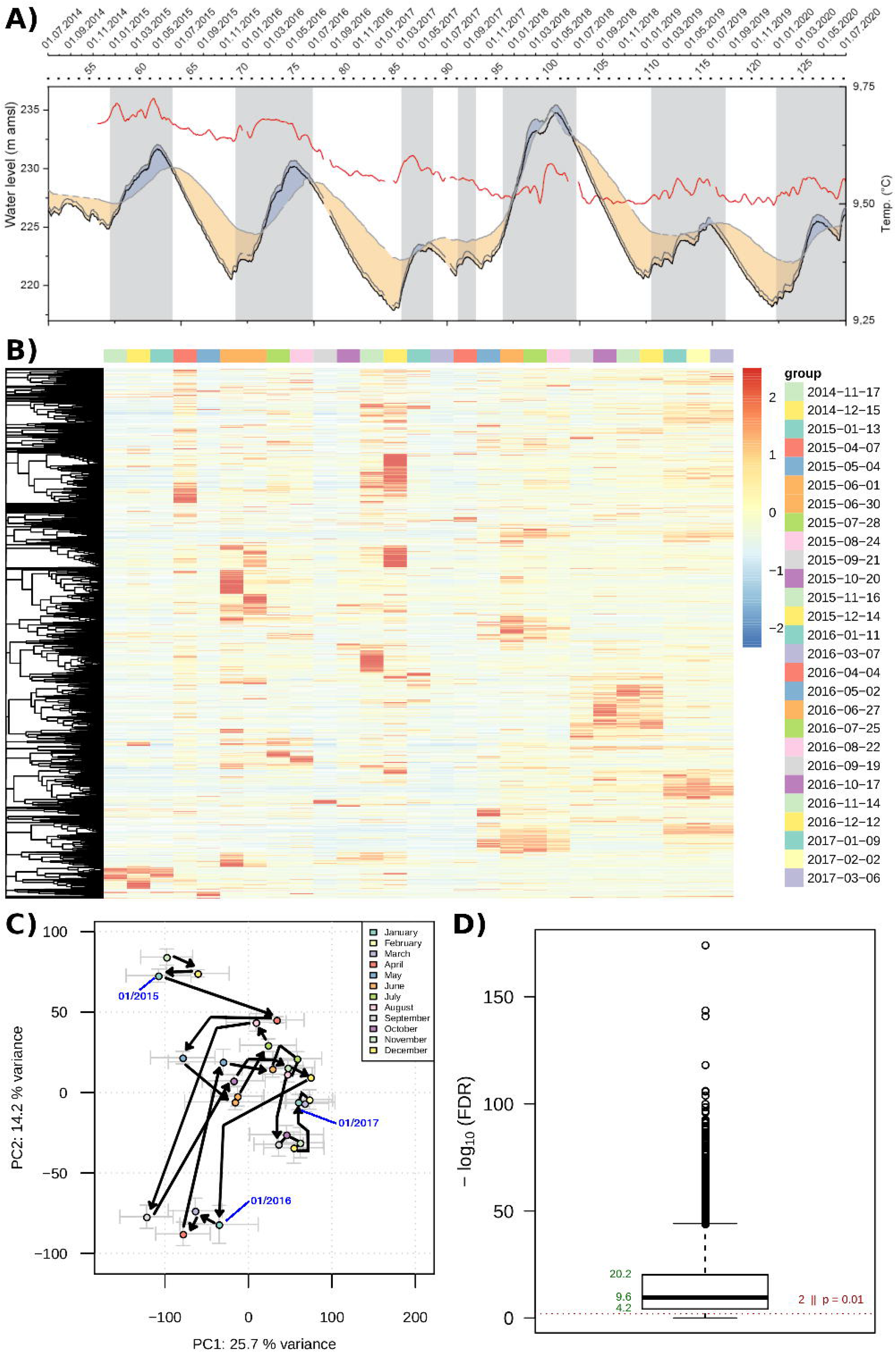
Single-well (H52) metabolome time-series analysis. **(A)** Groundwater level and temperature traces for H52 (blue; in comparison with H51 (black) and H53 (grey/yellow)). Recharge phases (grey-shaded) are inferred from the environmental tracer temperature (red) and multi-level head data. **(B)** Hierarchical cluster analysis of z-normalised metabolite feature intensities. **(C)** Principal component analysis of time-series. Arrows are drawn as guides for the chronological order of datapoints; the first datapoint of each year is labelled in blue font. **(D)** False discovery rate (FDR; from Benjamini-Hochberg corrected ANOVA) distribution as indicator of significance of metabolite feature intensity variability in the time series.

The metabolome in the phreatic groundwater (11/2014 to 03/ 2017) was also highly variable and exhibited singular emergences of metabolites at high intensities (Figure 3 B, D). Like in the well H32 in the uphill aeration zone and the other wells of the phreatic zone, recurrent fluctuations of metabolite signals were missing. Nevertheless, we again observed an association of the PCA with the groundwater fluctuation (Figure 3 A, C). The recharge-phase 11/2015-05/2016 contained a clear dispersion in the bottom-left of the PCA, while the subsequent discharge phase caused a cluster transition (08-09/2016, center-right) with another dispersion during the low-stand 01-03/2017 (centerright). While less distinct, the recharge phase 12/2014-05/2015 caused a PCA-shift (range 04-06/2015).

### General coincidence of temporal metabolome pattern changes and recharge dynamics across sampling sites

The association of metabolome PCAs with water levels emerged as a consistent observation across the recharge area. This was observed in shallow as well as in deep wells spanning different habitat conditions (Lazar et al., 2019) and microbial communities (Yan et al., 2020, 2021). In all cases (H32, H41 to −43, H51 to −53), the metabolome was variable, with pronounced shifts in the PCA-time-series, particularly in response to high-flow / recharge phases (Figures 2 and 3 for H32 and H52, as well as Supplementary Figures S3 to S8).

A summary is given in Supplementary Table 1, highlighting distinct phases in the PCAs. The association of the first two principal components with water level fluctuation, representing cross-stratal exchange in the fractured bedrock strata, revealed the strong responses to recharge dynamics. It is conceivable that high-flow phases will modify the DOM profile by co-transporting organic matter into the groundwater. In accordance, the dissolved organic carbon (DOC) levels in H32 and H52 were often variable during recharge periods (Supplementary Figures S1 and S2).

Transport of organic matter by water fluxes can change the groundwater metabolome by the input of novel compounds and microbial biochemical transformations will cause additional variability. At our study site, it has been recently reported that the microbiome composition is impacted by water recharge phases, as observed for the metabolome (Yan et al., 2020, 2021). It is conceivable that a changing microbiome will cause a shift in biochemical functions and therefore impact the metabolome composition. Other impacts on the metabolome will result from changing geochemical parameters, for instance, pH and oxygen levels that influence abiotic transformations.

The abovementioned processes can rationalize most of the metabolome profile variations over time. However, there remain unexplained changes. For instance, a disturbance in 09/2015 was observed in H52 (Figure 3), which was also observed in the other H5 wells (Supplementary Figures S7 and S8, Table S1), but not in the shallow H32 (Figure 2; information on location H4 not available). This disturbance could neither be explained by the sampling procedure (identical) nor did it have a clear representation in other measured parameters. A possible explanation might be that the corresponding pronounced recession period, followed by an extreme lasting groundwater highstand after 2013, might have influenced the prevalence of the specific metabolites.

### Inter-well metabolome similarity suggests cross-compartment exchange

Having established that water level fluctuations leave an imprint in the metabolome over time, we queried whether the metabolome also provides evidence for cross-compartment exchange along the hillslope recharge area (Figure 1). In addition to the metabolome data, we recorded quality parameters like temperature, dissolved oxygen, various solutes and DOC that have been previously used as environmental tracers to explore local aquifer systems (Figures S1 and S2) (Benischke, 2021; Jiang et al., 2021). These parameters fall short in resolution as aggregate parameters like DOC are too general, with only weak spatiotemporal variation. Also, major inorganic ions only allow interpretation at low resolution compared to the numerous organic analytes accessed by metabolomics techniques. Other tracers like fluorescent dyes or trace metals need to be deliberately released, thereby already impacting the environment. We, therefore, considered that untargeted LC/MS-based metabolomics monitors a suitable naturally-occurring organic tracer set for investigating cross-compartment exchange.

We considered metabolome data of the seven monitoring wells H32, H41 to - 43, and H51 to −53, covering the period from 11/2014 to 06/2020. For each timepoint, PCAs were generated to determine the principal discriminating components for inter-well dissimilarity (see Materials and Methods). From these, the PCA distances between every two wells were inferred and expressed as multiples of the distances’ standard deviations (Figure 4 and Supplementary Figures S9-S13). This allowed us to compare metabolome profiles acquired over the extended monitoring period.

**Figure 4.**
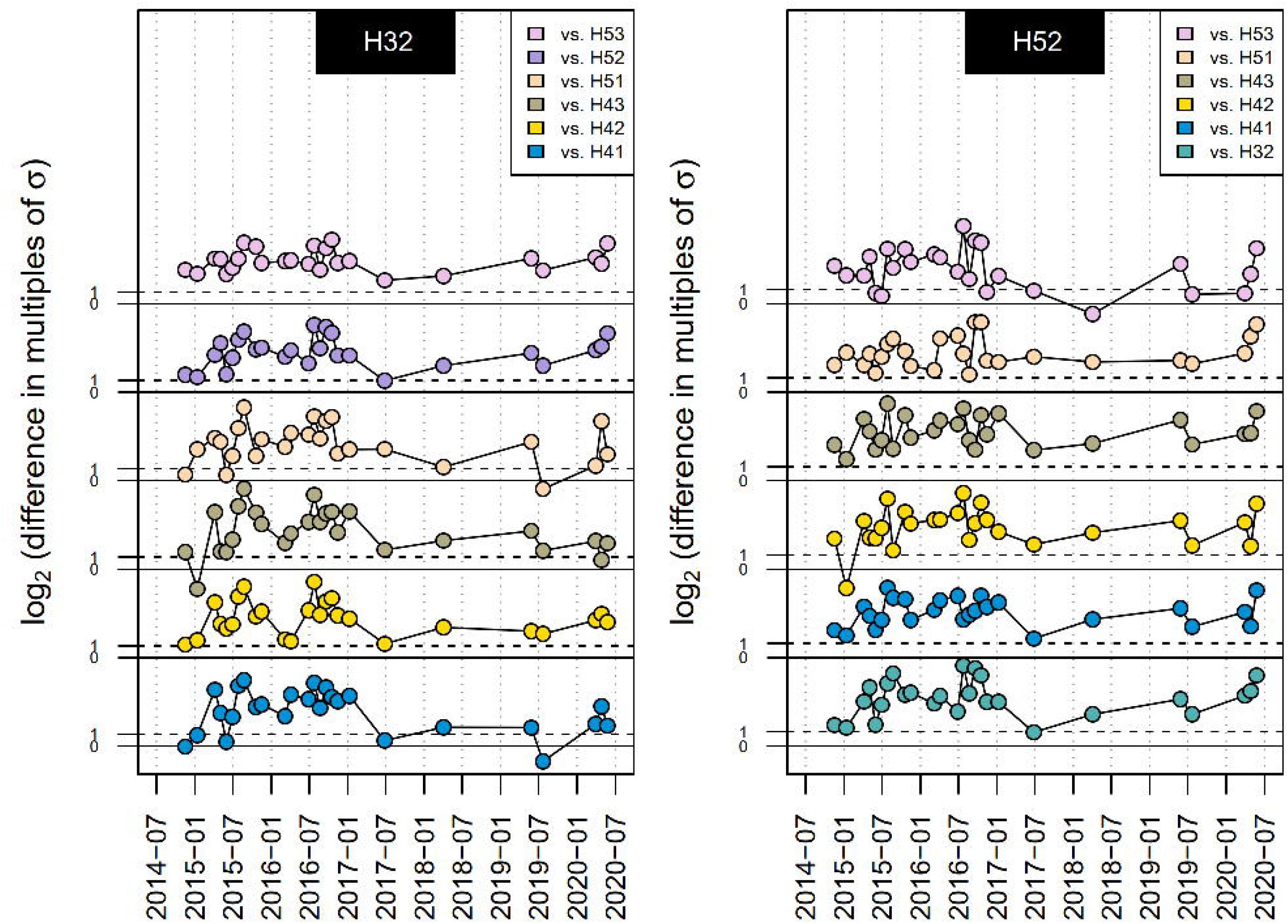
Metabolome dissimilarities of wells H32 (left) and H52 (right) compared against other wells included in the analysis (as shown in the legend). Well distances are expressed as multiples of the standard deviation σ in the PC1-PC2 space from principal component analysis (PCA) at individual sampling campaigns (i.e. time points). The distances were log_2_-transformed for presentation, and the 1σ and 2σ threshold are indicated (log_2_(1)=0 and log_2_(2)=1, respectively). Note for statistical assessment that for a normal-distributed random sample, the 1σ and 2σ ranges mark the intervals expectedly containing 68% and 95% of observations, respectively (Wang et al., 2015). Sampling campaign dates are detailed in Supplementary Table S2.

Well H52 was most distinct (i.e., had high scores for the dissimilarity) from the other wells (Figure 4); however, phases of similarity to well H53 emerged in 06-07/2015, 11/2016, 04/2018, 07/2019 and 04/2020. At all these timepoints, the H5 wells were at or near a peak of groundwater highstand (Figures 3 and Supplementary Figure S3), suggesting that ascending flow caused cross-stratal exchange and mixing. There were also phases of similarity between H52 and the well H51 at times around the highstands in 06/2015 and 09/2016, and an additional phase in 03/2016 where H51 and H52 were approaching their highstand.

Notably, in several phases when H52 and H53 were similar, well H51 had higher similarity to the aeration zone well H32 (04/2018; 2.5 km further upslope, see Figure 1) or the phreatic wells H41 to H-43 (07/2019 and 04/2020) than to the wells H52 and H53 (Supplementary Figure S13). These metabolome observations support earlier findings (Benk et al., 2019) of transient local flow patterns that connect even deep and isolated compartments with the surface or shallow upslope zones via ascending flow (R. Lehmann & Totsche, 2020).

In accordance, multivariate analyzes from aggregated, long-term Hainich monitoring data on the microbiome (Yan et al., 2020), physicochemical parameters (Kohlhepp et al., 2017; Schwab et al., 2017), and phospholipid-derived fatty acids (PLFA) (Schwab et al., 2017) classed the wells H52 and H53 as similar and distinct from the others, while a previous metabolomics study (Benk et al., 2019) placed H52 in proximity to H4 wells (1 km uphill, see Figure 1), but separate from H53 (same location, but 15 m above). The present analysis synergizes these findings by clarifying the changes in well-dissimilarities over longer monitoring periods.

The aeration zone well (H32), accessing a perched, oxic groundwater body, showed variable similarities with all wells but H53, which was consistently different. Phases of similarities to other wells were consistently observed following to all high-flow phases in H32, as follows: 11/2014-01/2015 (to H41-43, H51-52), 06/2015 (H41, H43, H51-52), 03-03/2016 (H42), 07/2017 (H41-43, H52), 04/2018 (H41, H51), 07/2019 (H41, H43, H51) and 04-05/2020 (H41, H43, H51).

This demonstrates that PCA of metabolomics data from long term surveys can be used to derive time-traces of dissimilarities by inferring the sample distances in the PC1-PC2 space. Acknowledging the obstacles of long-term data acquisition, where metabolomics measurement parameters, such as operator, device calibration, separation column, and method may have changed, the extraction of PCA-distances per time point allows integration of long-term monitoring data. We can conclude that the sedimentary-rock groundwaters vary in dissimilarity over time across the hillslope recharge area. Phases of low dissimilarity emerge coincidently with groundwater recharge dynamics, covering surface-fed as well as subsurface matter exchange.

Previously, it has been queried to which extent the surface vegetation or (agricultural) land use in a recharge area impacts underlying groundwater systems (Küsel et al., 2016). Metabolome investigations of soil waters sampled in lysimeters revealed discrete profiles for the input metabolome in forest, pasture and agricultural areas (Ueberschaar et al., 2021). For the groundwater wells in Figure 1, we have demonstrated elsewhere that the metabolomes of the wells studied on a single sampling day were discernible in relation to the factors land-use type, aquifer system and sampling depth (Sanchez-Arcos et al., 2022). We can now show that for the long-term groundwater metabolomes (Figure 4), vertical transport or the locations’ respective land use types on the surface are not the major determinants. Instead, periods of similarity occur between H32 and wells at the H4 and H5 sites with different land use and different aquifer properties (Lazar et al., 2019; R. Lehmann & Totsche, 2020). No continuous segregation by land-use type was observed, implying that its impact on the well metabolomes is either transient or secondary. Changing similarities are meanwhile mainly driven by variable flow patterns and water stands. Notably, these may become more fluctuating with weather and climatic extremes, both which are expected to become more frequent under climate change predictions (IPCC, 2022). In effect, it can be expected that groundwater metabolic diversity may rise further thus posing unprecedented challenges to groundwater microbes and ecosystems.

## Conclusion

We show that groundwater metabolomics can resolve the sum parameter DOC with much detail and disclose hidden information on groundwater ecosystems’ functioning and dynamics. We further demonstrate that groundwater flows exert a major impact on the groundwater metabolome variability, which will thus affect the carbon and energy provision to belowground microbial communities. Such information is essential to discover ecosystem patterns and responses, such as groundwater recharge events or climatic extremes that are expected to increase in severity with climatic extremes predicted in climate change scenarios and may impact the discharge-recharge cycle. Generally, the groundwater metabolome and its temporal dynamics provide cross-habitat information from multiple sources of metabolites, covering contributions from microbial populations, as well as abiotic transformations. This allows, for example, the recognition of similarities caused by high-flow phases in wells that are several kilometers apart. The metabolomics approach has the advantage over tracer experiments that it determines natural organic components of the groundwater without the need to introduce markers. It closes a critical gap in the interdisciplinary investigation of groundwater ecosystems.

## Acknowledgments

This project was funded by the Collaborative Research Centre AquaDiva of the Deutsche Forschungsgemeinschaft (DFG, German Research foundation – SFB 1076 Project number 218627073). We thank Falko Guthmann and Heiko Minkmar for maintaining the Hainich CZE infrastructure and conducting sampling, and Tim Baumeister and Ann-Sophie Lehnert for support in sample processing.

## Data and materials availability

Data acquired by liquid chromatography coupled to mass spectrometry is accessible via the Metabolights repository (Haug et al., 2019) https://www.ebi.ac.uk/metabolights with study identifier MTBLS3450.

https://www.ebi.ac.uk/metabolights/reviewer13c7b37d-1178-421b-93d5-f19c06ea545c (Provisional Review Link)

