## Supplementary material for "Groundwater metabolome responds to recharge in fractured sedimentary strata": Figure S2

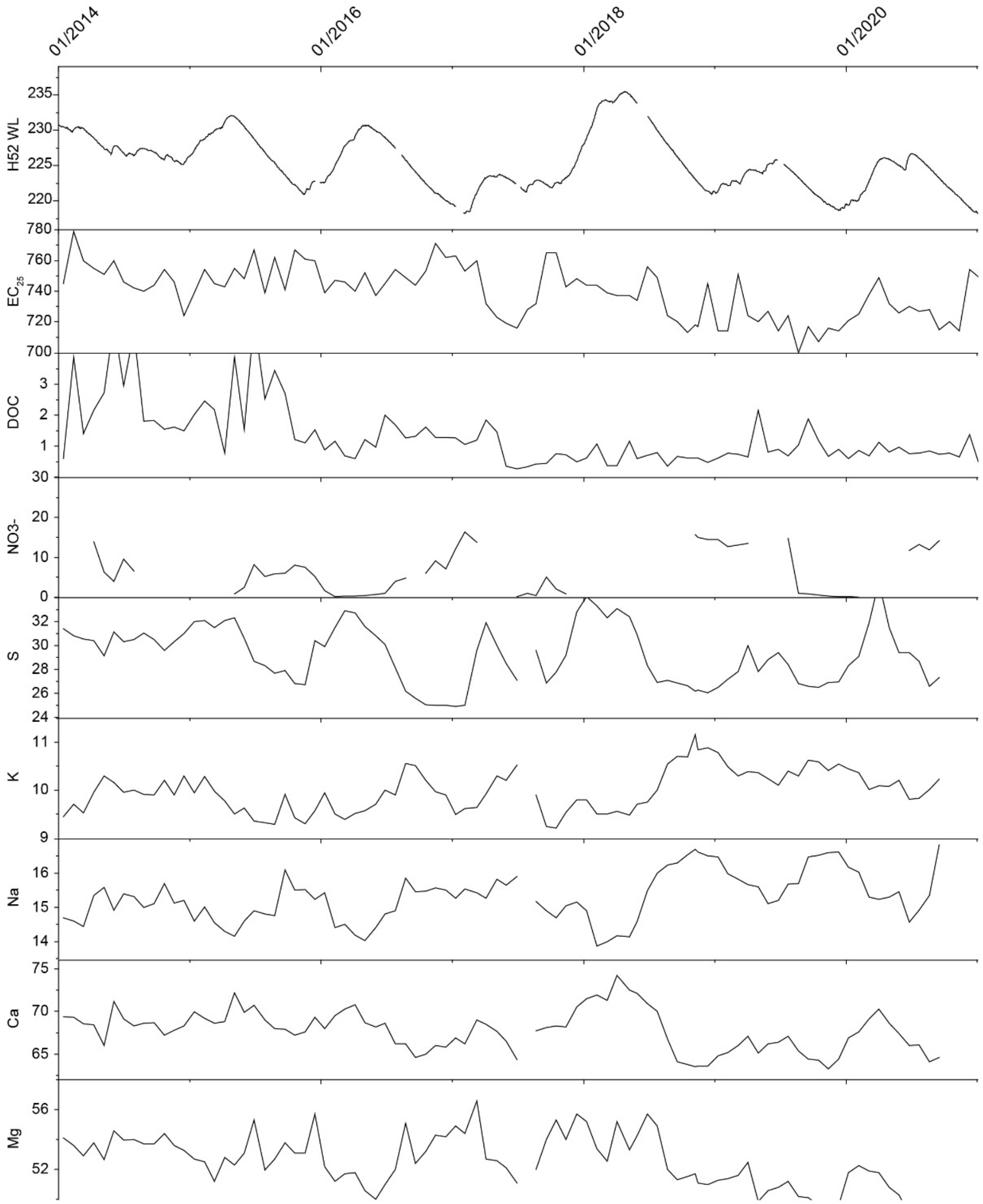


**Figure S2.** Fluctuation of groundwater level and quality in the phreatic zone well (H52): water level (WL), conductivity ($EC_{25}$), dissolved organic carbon (DOC), and concentrations of various elements / ions (after filtration 0.45 µm, S for sulfur in any chemical composition).
