## Supplementary material for "Groundwater metabolome responds to recharge in fractured sedimentary strata": Figure S3

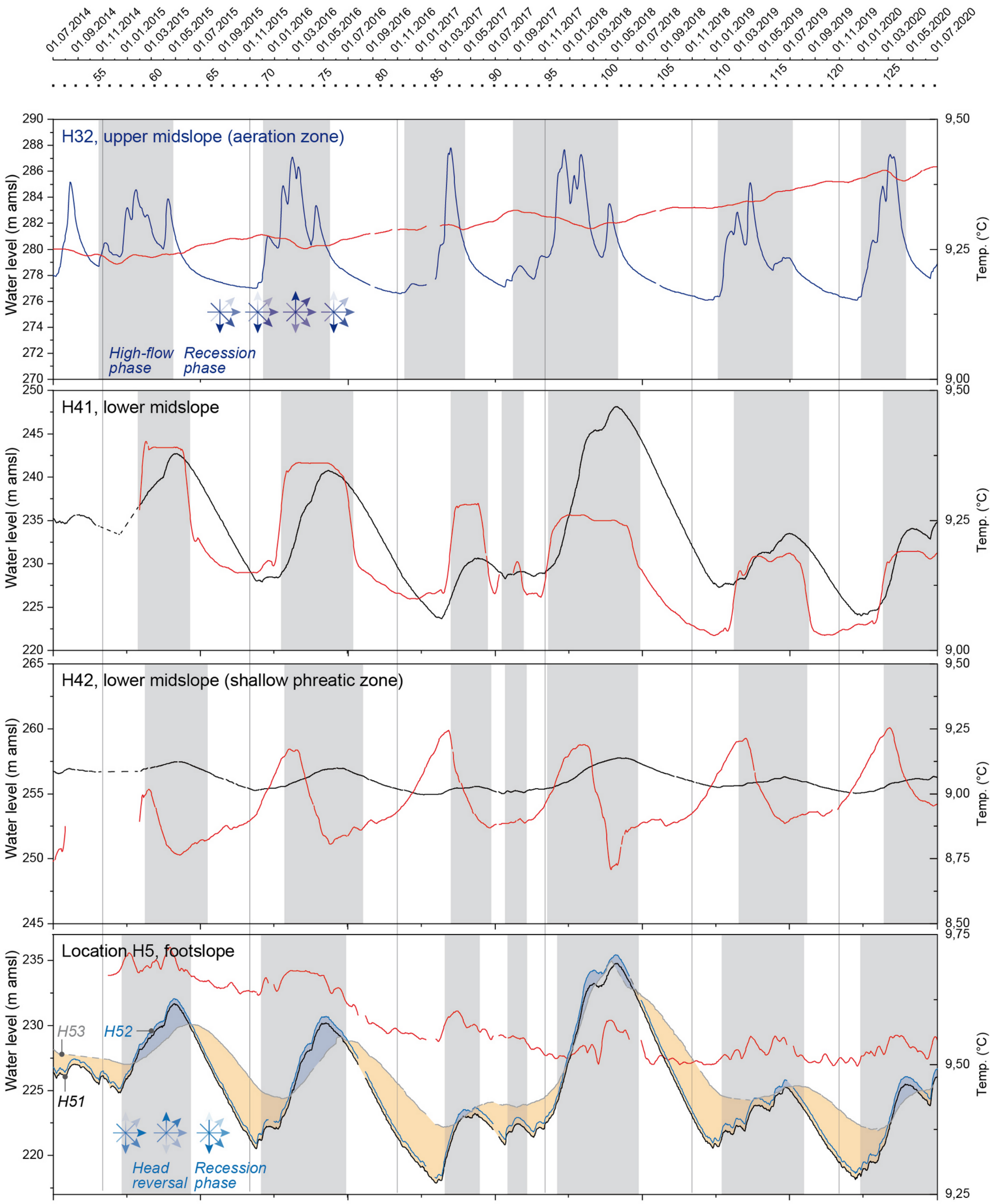


**Figure S3.** Time series of groundwater levels and temperatures along the topographic recharge area. Campaign numbers are shown on top. Contrasting seasonal phases (grey-shaded) are inferred from the environmental tracer temperature (red) and multi-level head data (H5). Arrow roses mark presumed directions of transient cross-stratal flow and matter exchange.
