## Supplementary material for "Groundwater metabolome responds to recharge in fractured sedimentary strata": Figure S6

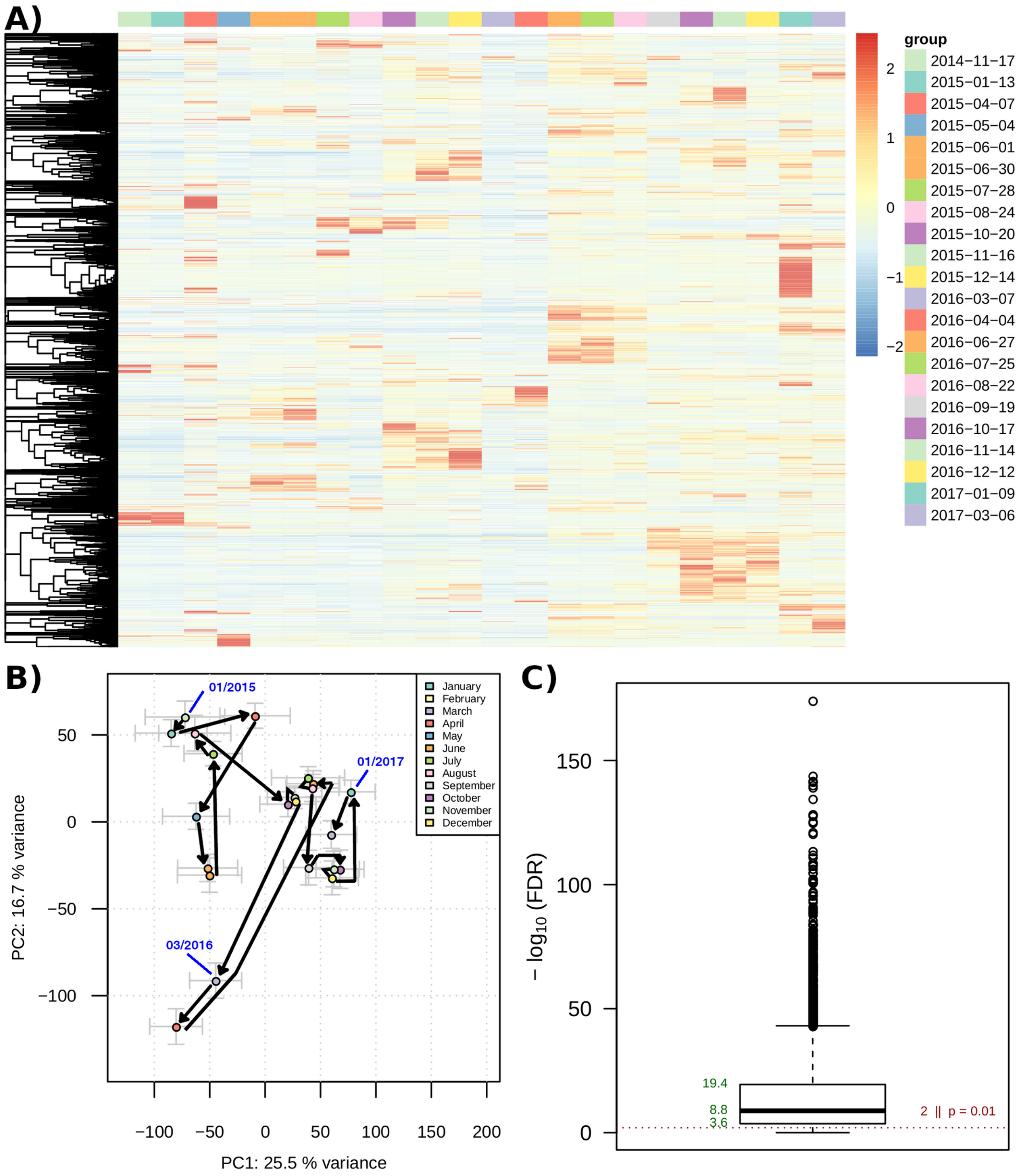


**Figure S6.** Metabolome time-series analysis of the shallow phreatic zone at the lower midslope (well H43). (A) Hierarchical cluster analysis of z-normalised metabolite feature intensitie (values with $z > 2.5$ and $z < -2.5$ not further distinguished by colour). (B) Principal component analysis of time-series. Arrows are drawn as guides for chronological order of datapoints. (C) False discovery rate (FDR; from Benjamini-Hochberg corrected ANOVA) distribution as indicator of significance of metabolite feature intensity variability in the time series.
