## Supplementary material for "Groundwater metabolome responds to recharge in fractured sedimentary strata": Figure S9

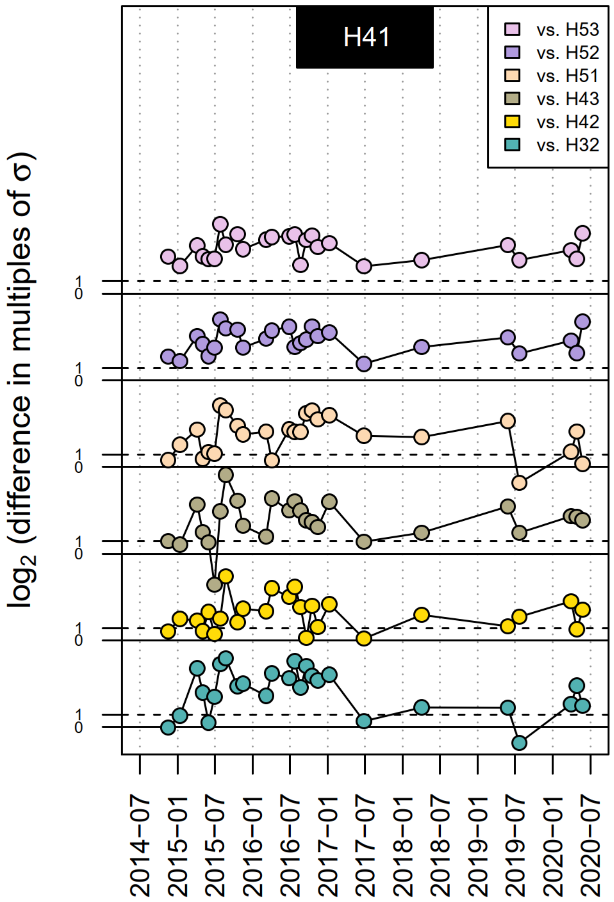


**Figure S9.** Dissimilarity between the H41 metabolome and the other monitoring wells. Sampling campaigns included in this analysis are detailed in Supplementary Table S1.
