## Supplementary material for "Groundwater metabolome responds to recharge in fractured sedimentary strata": Figure S10

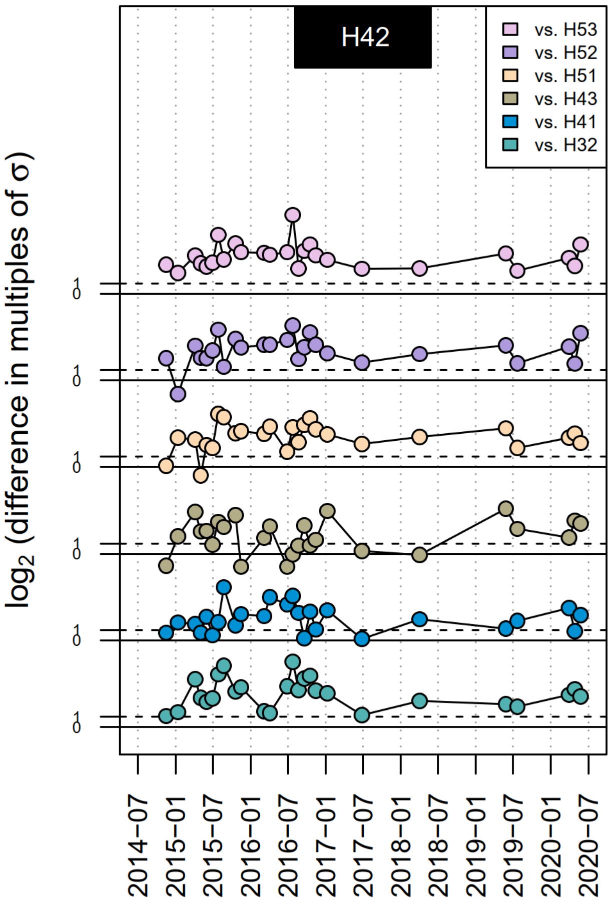


**Figure S10.** Dissimilarity between the H42 metabolome and the other monitoring wells. Sampling campaigns included in this analysis are detailed in Supplementary Table S1.
