## Supplementary material for "Groundwater metabolome responds to recharge in fractured sedimentary strata": Figure S12

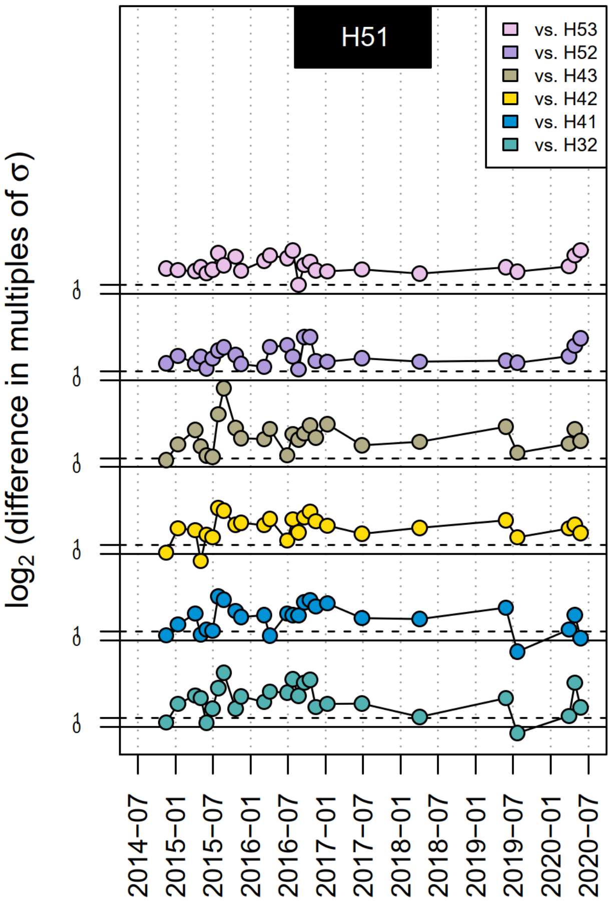


**Figure S12.** Dissimilarity between the H51 metabolome and the other monitoring wells. Sampling campaigns included in this analysis are detailed in Supplementary Table S1.
