## Supplementary material for "Groundwater metabolome responds to recharge in fractured sedimentary strata": Table S1

| Sampling campaign date | H32 | H41 | H42 | H43 | H51 | H52 | H53 |
|---|---|---|---|---|---|---|---|
| 2014-09-23 | | | | | | | |
| 2014-11-17 | 1 | | 1 (high variability phase) | 1.1 | 1 | 1 | |
| 2014-12-15 | 1 | | 1 (high variability phase) | 1.1 | 1 | 1 | 1.1 |
| 2015-01-13 | 1 | 1.1 | 1 (high variability phase) | 1.1 | 1 | 1 | 1.1 |
| 2015-04-07 | 2 | 1.1 | 1 (high variability phase) | 1.1 | 2 (high variability phase) | 2 (high variability phase) | 1.1 |
| 2015-05-04 | 2 | 2 | 1 (high variability phase) | 2 | 2 (high variability phase) | 2 (high variability phase) | 1.1 |
| 2015-06-01 | 2 | 2 | 1 (high variability phase) | 2 | 2 (high variability phase) | 2 (high variability phase) | 1.1 |
| 2015-06-30 | 2 | 2 | 1 (high variability phase) | 2 | 2 (high variability phase) | 2 (high variability phase) | 1.1 |
| 2015-07-28 | 2 | 1.2 | 1 (high variability phase) | 1.2 | 2 (high variability phase) | 2 (high variability phase) | 1.1 |
| 2015-08-24 | 2 | 1.2 | 1 (high variability phase) | 1.2 | 2 (high variability phase) | 2 (high variability phase) | 1.1 |
| 2015-09-21 | 2 | | 1 (high variability phase) | | 3 | 3 | 2 |
| 2015-10-20 | 2 | 3.1 | 1 (high variability phase) | 3.1 | 4 | 4.1 | 1.2 |
| 2015-11-16 | 2 | 3.1 | 1 (high variability phase) | 3.1 | 4 | 4.1 | 1.2 |
| 2015-12-14 | 2 | 3.1 | 1 (high variability phase) | 3.1 | 4 | 4.1 | 1.2 |
| 2016-01-11 | 3 | | | | 5 | 5 | 3 |
| 2016-03-07 | 3 | 4 | 2 | 4 | 5 | 5 | 3 |
| 2016-04-04 | 3 | 4 | 2 | 4 | 5 | 5 | 3 |
| 2016-05-02 | | 3.2 | 3 | | 6 | 4.2 | |
| 2016-06-27 | 4 | 3.2 | 3 | 3.2 | 6 | 4.2 | 4 (high variability phase) |
| 2016-07-25 | 4 | 3.2 | 3 | 3.2 | 6 | 4.2 | 4 (high variability phase) |
| 2016-08-22 | 4 | 3.2 | 3 | 3.2 | 6 | 4.2 | 4 (high variability phase) |
| 2016-09-19 | 4 | 5 | 4 | 5 | 7 | 6 | 5 |
| 2016-10-17 | 4 | 5 | 4 | 5 | 7 | 6 | 5 |
| 2016-11-14 | 4 | 5 | 4 | 5 | 7 | 6 | 5 |
| 2016-12-12 | 4 | 5 | 4 | 5 | 7 | 6 | 5 |
| 2017-01-09 | 4 | 6 | 5 | 6 | 8 | 7 | 5 |
| 2017-02-02 | 4 | 6 | 5 | 6 | 8 | 7 | 5 |
| 2017-03-06 | 4 | 6 | 5 | 6 | | 7 | 5 |

**Table S1.** Overview table of emerging clusters and disturbances in time-series PCAs (Figures 1-7, panels B). For each well, black fields indicate dates in which a sample from the well was drawn. The coloured cells mark phases based on proximity of PCA datapoints. Colours and numbers distinguish subsequent phases for each well only, not suggesting shared phases or relationships between the wells.
