## Supplementary material for "Groundwater metabolome responds to recharge in fractured sedimentary strata": Table S2

| **Joint Sampling Campaign #** | **Date (first day of campaign)** |
|---|---|
| 56 | 2014-11-17 |
| 58 | 2015-01-13 |
| 61 | 2015-04-07 |
| 62 | 2015-05-04 |
| 63 | 2015-06-01 |
| 64 | 2015-06-30 |
| 65 | 2015-07-28 |
| 66 | 2015-08-24 |
| 68 | 2015-10-20 |
| 69 | 2015-11-16 |
| 73 | 2016-03-07 |
| 74 | 2016-04-04 |
| 77 | 2016-06-27 |
| 78 | 2016-07-25 |
| 79 | 2016-08-22 |
| 80 | 2016-09-19 |
| 81 | 2016-10-17 |
| 82 | 2016-11-14 |
| 84 | 2017-01-09 |
| 90 | 2017-06-26 |
| 100 | 2018-04-03 |
| 115 | 2019-05-27 |
| 117 | 2019-07-22 |
| 126 | 2020-03-30 |
| 127 | 2020-04-27 |
| 128 | 2020-05-25 |

**Table S2.** Dates of sampling campaigns included in the well-dissimilarity time-series.
